# Conditional Stomatal Closure in a Fern Shares Molecular Features with Flowering Plant Active Stomatal Responses

**DOI:** 10.1101/2021.03.06.434194

**Authors:** Andrew R.G. Plackett, David M. Emms, Steven Kelly, Alistair M. Hetherington, Jane A. Langdale

## Abstract

Stomata evolved as plants transitioned from water to land, enabling carbon dioxide uptake and water loss to be controlled. In flowering plants, the most recently divergent land plant lineage, stomatal pores actively close in response to drought. In this response, the phytohormone abscisic acid (ABA) triggers signalling cascades that lead to ion and water loss in the guard cells of the stomatal complex, causing a reduction in turgor and pore closure. Whether this stimulus-response coupling pathway acts in other major land plant lineages is unclear, with some investigations reporting that stomatal closure involves ABA but others concluding that closure is passive. Here we show that in the model fern *Ceratopteris richardii* active stomatal closure is conditional on sensitisation by pre-exposure to either low humidity or exogenous ABA and is promoted by ABA. RNA-seq analysis and *de novo* transcriptome assembly reconstructed the protein coding complement of the *C. richardii* genome with coverage comparable to other plant models, enabling transcriptional signatures of stomatal sensitisation and closure to be identified. In both cases, changes in abundance of homologs of ABA, Ca^2+^ and ROS-related signalling components were observed, suggesting that the closure response pathway is conserved in ferns and flowering plants. These signatures further suggested that sensitisation is achieved by lowering the threshold required for a subsequent closure-inducing signal to trigger a response. We conclude that the canonical signalling network for active stomatal closure functioned in at least a rudimentary form in the stomata of the last common ancestor of ferns and flowering plants.

**Significance Statement:** Stomata are valve-like pores that control the uptake of CO_2_ and the loss of water vapour in almost all land plants. In flowering plants, stomatal opening and closure is actively regulated by a stimulus-response coupling network. Whether active stomatal responses are present in other land plant lineages such as ferns has been hotly debated. Here we show that stomatal responses in the fern *Ceratopteris richardii* are active but depend on their past growth environment, and demonstrate that fern stomatal closure and sensitisation are associated with the altered expression of genes whose homologs function in the canonical stomatal regulatory network of flowering plants. Genetic pathways for active stomatal regulation therefore most likely evolved before the divergence of ferns and flowering plants.

## Introduction

Stomata are pores present on the surfaces of plant leaves that control the uptake of CO_2_ and the loss of water vapour. The acquisition of stomata was a key land plant adaptation, which together with the development of a waxy cuticle and vascular system allowed early plants to colonise the terrestrial environment[^1^]. The ability to control CO_2_ uptake is important in the context of photosynthesis, whereas the regulation of evapotranspiratory water loss impacts on water and mineral nutrient accumulation in the aerial parts of the plant, protects against short periods of reduced soil water availability and provides leaf cooling capacity. In addition, stomatal closure provides protection against invasion by some pathogens. The regulation of stomatal aperture by light, humidity, atmospheric CO_2_, and the plant hormone abscisic acid (ABA) has been extensively investigated in the model flowering plant *Arabidopsis thaliana*. These studies identified networks of intracellular signalling proteins and second messengers that act in the guard cells of the stomatal complex, to couple extracellular stimuli to an opening or closing response[^2,3,4^]. Although stomata are present in all land plant lineages except liverworts[^5^], in which they have likely been secondarily lost[^6^], the question of when active stomatal closure mechanisms evolved remains hotly debated.

Studies of stomatal responses in non-flowering plant species have led to conflicting interpretations of underlying mechanisms. Observations that stomata close in response to ABA and CO_2_ in two moss species[^7^], *in silico* evidence suggesting that stomata evolved only once[^6^] and the presence of genes encoding components of the *Arabidopsis* stomatal closure pathway in moss genomes[^8,9^] could indicate that active stomatal responses are ancient. However, reports from non-flowering vascular plants have been contradictory. Some suggest that in lycophytes and ferns stomatal closure is hydropassive and guard cells are insensitive to closure-inducing signals such as ABA and high CO_2_ levels[^10,11,12,13,14^] whilst in contrasting reports stomata in the lycophyte *Selaginella uncinata*[^15^] and a number of ferns[^16,9,17, 18,19, 20^] were shown to close in response to ABA and/or CO_2_. The observation that stomatal closure in response to exogenous ABA is conditional on growth conditions in some (but not all) species of fern[^17^] may highlight the confounding variable in previous reports, but the underlying molecular basis for fern stomatal closure and conditional responsiveness is unknown.

The application of modern genetic analysis approaches to the fern lineage remains challenging, with few techniques and resources available. To identify the pathways underlying fern stomatal closure mechanisms, we utilised RNA-seq technology and *de novo* transcript assembly to generate and compare transcriptome profiles associated with different stomatal responses in the fern *Ceratopteris richardii*. It has previously been suggested that stomatal closure in *C. richardii* is not activated by the canonical ABA-induced response pathway found in flowering plants, based on the observation that stomatal conductance is reduced in response to increased water vapour deficit even in a mutant lacking a functional homolog of a receptor kinase required for ABA-induced stomatal closure in *Arabidopsis*[^21^]. However, *C. richardii* is a neotropical semi-aquatic fern that is routinely grown at high humidity (i.e. low water vapour deficit) in the laboratory[^22,23^], and stomatal responses may have been masked by conditional behaviours now known from other ferns[^17^]. We demonstrate herein that the stomata of *C. richardii* can actively close in response to both low humidity and ABA but that this response is dependent on previous sensitisation by either low humidity or ABA. Our RNA-seq analysis provides evidence that the signalling network controlling stomatal closure in ferns contains homologs of ABA transport, Ca^2+^ signalling and ROS signalling components from canonical *Arabidopsis* stomatal closure pathways, and suggests that sensitisation potentiates these same pathways to respond to closure stimuli at a lower threshold. We conclude that a core of regulatory networks that control active stomatal closure is conserved between ferns and flowering plants.

## Results

### *Active stomatal closure in* C. richardii *is conditional on pre-sensitisation by low humidity or exogenous ABA*

To test whether active stomatal closure in *C. richardii* is conditional on growth environment, stomatal responses were compared between two humidity pretreatments. Plants grown either in constant high relative humidity (94.9 ± 0.36%) (referred to hereafter as wet-grown) or periodically exposed to low (ambient) humidity (48.3 ± 0.69%) (referred to hereafter as dry-pretreated) for ten minutes (see Materials and Methods). After an interval of growth at high humidity to ensure all stomatal were open, both pretreatment groups were exposed to low humidity for 120 minutes. Stomata of fronds from dry-pretreated plants exhibited a significant (p < 0.05) and progressive reduction in stomatal pore area in response to low humidity at both 60 and 120 minutes (**Fig. 1*A***), with mean pore areas of 79.7% and 54.0% of the mean area measured at 0 minutes, respectively (**Dataset S1**). Simultaneous application of exogenous ABA significantly enhanced this response (p < 0.05) at both 60 (66.9%) and 120 minutes (40.0%) (**Fig. 1*A***). In contrast, stomata of wet-grown plants showed a significant reduction in pore area (p < 0.05) only after 120 minutes exposure to low humidity and of much lesser magnitude than in dry-pretreated plants (89.0% of mean pore area at 0 minutes), with no response to ABA (p > 0.05) (**Fig. 1*A***). Active stomatal closure in dry-pretreated plants is further supported by a specific reduction in pore aperture width and not length (**SI Appendix**, **Fig. S1**, **S2**), and with a greater resilience to wilting than seen in wet-grown plants (**SI Appendix**, **Fig. S3**), with no change in stomatal density (**SI Appendix**, **Fig. S1**). When assays were repeated under constant high humidity the responses of stomata from dry-pretreated plants were much reduced, with mean pore area at 60 and 120 minutes 81.6% and 76.2% that of 0 minutes respectively, no significant change between 60 and 120 minutes (p > 0.05) (**Fig.1*B***) and no enhancement of closure by exogenous ABA (**Fig. 1*B***). Stomatal responses of wet-grown plants were similar between 120 minutes constant high humidity (86.1% of mean pore area at 0 minutes) (**Fig. 1*B***) and low humidity. Dry pretreatment was thus necessary to sensitise stomata for active closure in response to a low humidity stimulus, and low humidity was required to see any effect of an ABA stimulus on closure.

**Figure 1.**
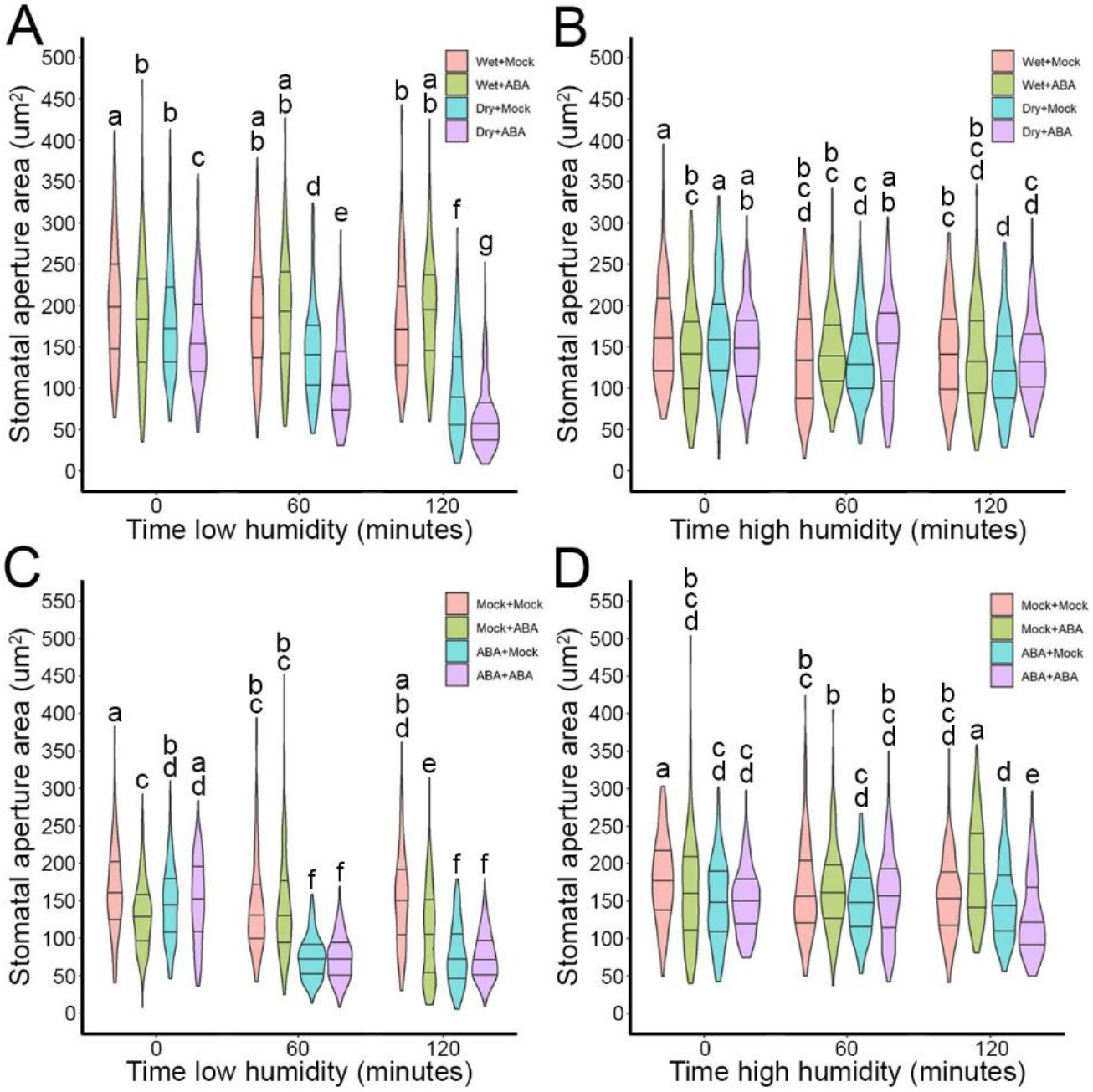
Active closure of *C. richardii* stomata in response to low humidity and ABA is conditional on pre-exposure to either stimulus. **A-D.** The response of stomatal aperture areas on fronds of wet-grown (‘Wet’) or dry-pretreated (‘Dry’) plants (**A,B**) or wet-grown plants pretreated with either a mock (‘Mock’) or 100uM exogenous ABA (‘ABA’) solution (**C,D**) exposed to a prolonged low humidity stimulus (**A,C**) or maintained in high humidity conditions (**B**,**D**) for 0, 60 or 120 minutes (as shown). Either mock solution (‘+Mock’) or 100uM exogenous ABA (+ABA) was applied in an orthogonal design at the start of the assay. *N* = 180 stomata per pretreatment + treatment combination, measured from three independent plants per timepoint. Violin plots represent data distribution density, with thickness corresponding to frequency of datapoints at that value. Median value and upper and lower quartiles are represented by the middle, upper and lower horizontal lines, respectively. Pairwise comparisons were performed using two-tailed Mann-Whitney tests. Where necessary to meet the assumptions of this test, comparisons were performed on square root-transformed datasets (**A**,**C**,**D**). Letters denote statistically significant differences (p < 0.05) within each plot. Pre-treatment + treatment combinations with the same letter are not significantly different from each other. Individual stomatal measurements are provided in **Dataset S1**.

To test whether stomata could also be sensitised solely by ABA, wet-grown plants were pretreated with periodic application of exogenous ABA or a mock control solution (see Materials and Methods). After subsequent exposure to a low humidity stimulus, stomata from ABA-pretreated plants showed a significant reduction in stomatal area compared to mock-pretreated controls (p < 0.05) at 60 and 120 minutes (**Fig. 1*C***), achieving the same magnitude of response by 60 minutes (51.1% of mean pore area at 0 minutes) as seen in dry-pretreated plant by 120 minutes (**Dataset S1**). Exogenous ABA treatment in addition to the low humidity stimulus did not enhance this response (p > 0.05, **Fig. 1*C***), presumably because maximum closure had already been reached by 60 minutes. Stomata from mock-pretreated plants were essentially unresponsive to low humidity, with a transient reduction in aperture area at 60 minutes only (p < 0.05) to 87.7% of mean pore area at 0 minutes (**Fig. 1*C***). As with dry-pretreated plants, ABA-pretreated plants showed greater resistance to wilting compared with controls (**SI Appendix**, **Fig. S3**), with no change in stomatal density (**SI Appendix**, **Fig. S1**). When assayed under high humidity the closure responses of stomata from ABA-pretreated plants were lost (p > 0.05, **Fig. 1*D***), but exogenous ABA did elicit a minor closure response (p < 0.05) after 120 minutes (87.0%) (**Fig. 1*D***; **Dataset S1**). Stomata from mock-pretreated plants exhibited a similar degree of responsiveness under both low and high humidity (**Fig. 1*C*,*D***; **Dataset S1**). Together these results show *C. richardii* stomata can be sensitised by ABA without low humidity, enabling subsequent active closure in response to either low humidity or ABA.

### *Homologs of* Arabidopsis *regulators of stomatal aperture and ABA responses are differentially expressed during stomatal sensitisation in* C. richardii

To identify genetic components of the mechanisms underlying stomatal sensitisation in *C. richardii*, RNA-seq was used to establish genome-wide transcript profiles in fronds of wet-grown, dry-pretreated, and ABA-pretreated plants (**Fig. 2*A***). The completeness of the predicted proteome contained within the *de novo* transcriptome assembled was similar to that of complete plant genomes and substantially more complete than the published *C. richardii* partial genome assembly[^24^] (**Fig. 2*B***). The majority of protein-coding transcripts (68.7%) had identifiable homology to *Arabidopsis* genes (see Materials and Methods), of which 49% were direct orthologs to *Arabidopsis* genes (**Fig. 2*C***). The quality of our *de novo* proteome was thus sufficient to identify conserved genetic networks between *Arabidopsis* and *C. richardii*.

**Figure 2.**
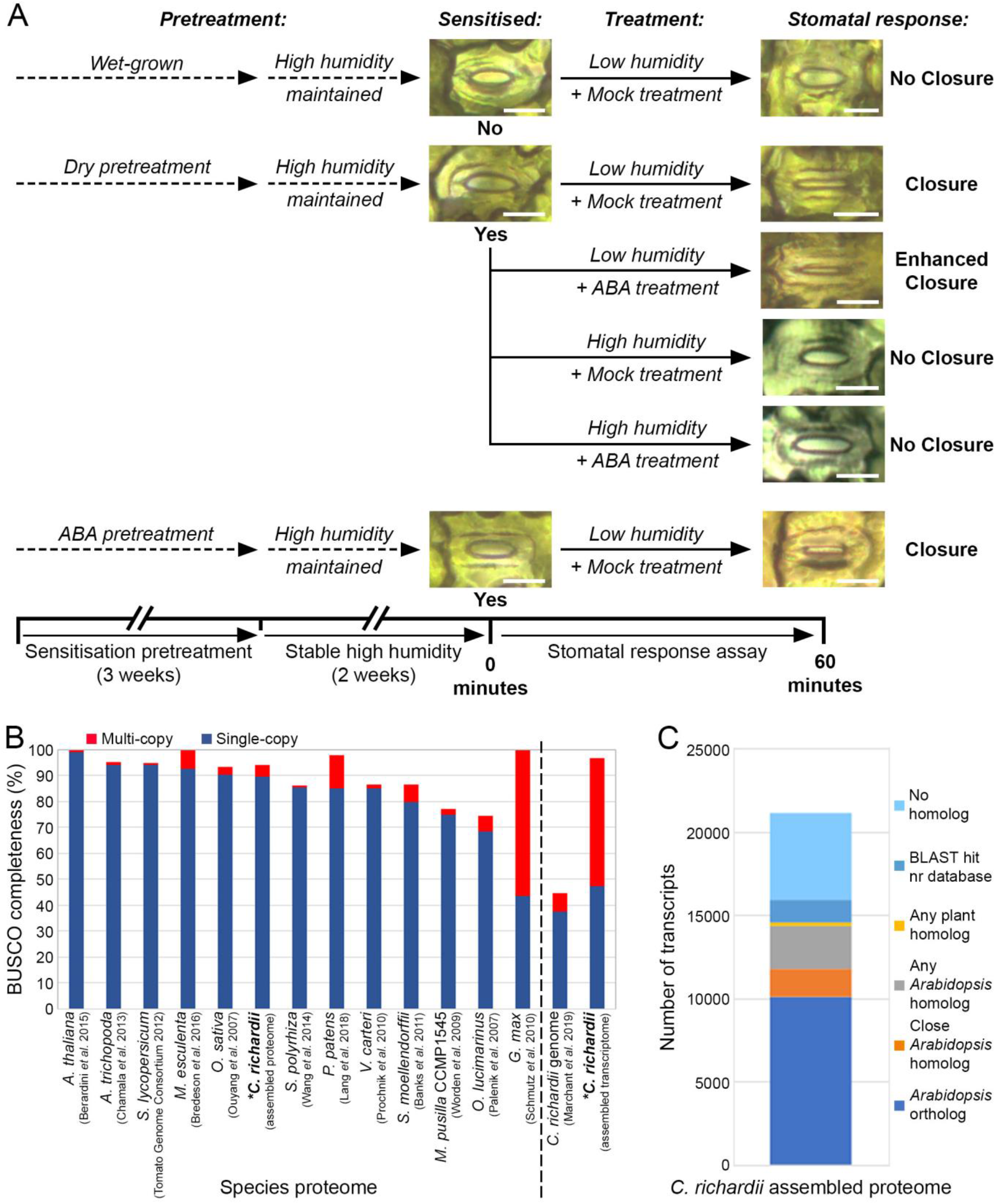
*De novo* transcriptome assembly provides a robust platform for the identification of stomatal sensitisation and closure signatures. **A.** Sampling for RNA-seq analysis followed the design of stomatal response assays to capture whole-frond transcriptomes at 0 or 60 minutes after humidity stimulus under selected pretreatment and treatment combinations, as shown. The corresponding stomatal aperture status for each timepoint is shown, as is the observed stomatal sensitivity and closure response. Scale bars = 25μm. **B.** Benchmarking Universal Single-Copy Orthologue (BUSCO) analysis of transcript completeness within the *C. richardii do novo* transcript population assembled from (**A**). Protein-coding transcripts (proteome) were compared against equivalent proteomes from reference species in the OrthoFinder analysis used to assign transcript homology (see Materials and Methods), and all *de novo* assembled *C. richardii* transcripts were compared against the published genome assembly[^24^]. The proteome assembled here is of equivalent completeness to reference proteomes in other land plant species. Asterisks denote datasets generated by this study. **C**. The degree of conservation and the extent of homology between *C. richardii* protein-coding transcripts identified within the *de novo* assembly and the *Arabidopsis* annotated genome[^61^].

A comparison of transcriptome profiles in wet-grown (non-sensitised) versus dry-pretreated or ABA-pretreated (sensitised) fronds revealed 67 transcripts with abundance levels that were significantly different (p < 0.05) in wet-grown versus dry-pretreated fronds (**Fig. 3*A***) and 6919 transcripts with significantly different (p < 0.05) levels in wet-grown and ABA-pretreated fronds (**Fig. 3*A***; **SI Appendix, Fig. S4**). The large number of genes responding to ABA pretreatment was anticipated because there were pleiotropic effects on frond morphology (**SI Appendix**, **Fig. S1**) and ABA is known to impact on other processes that are unrelated to stomatal physiology. However, just 47 transcripts displayed significant differences in abundance (42 at p<0.01; 5 at 0.01<p<0.05) in both wet-grown versus dry-pretreated and wet-grown versus ABA-pretreated comparisons (**Fig. 3*A***). These 47 transcripts, 11 of which had significantly higher levels in sensitised fronds and 36 of which had lower levels (p < 0.05) (**Fig. 3*B***; **Dataset S2**), thus represent the stomatal sensitisation signature in *C. richardii*.

**Figure 3.**
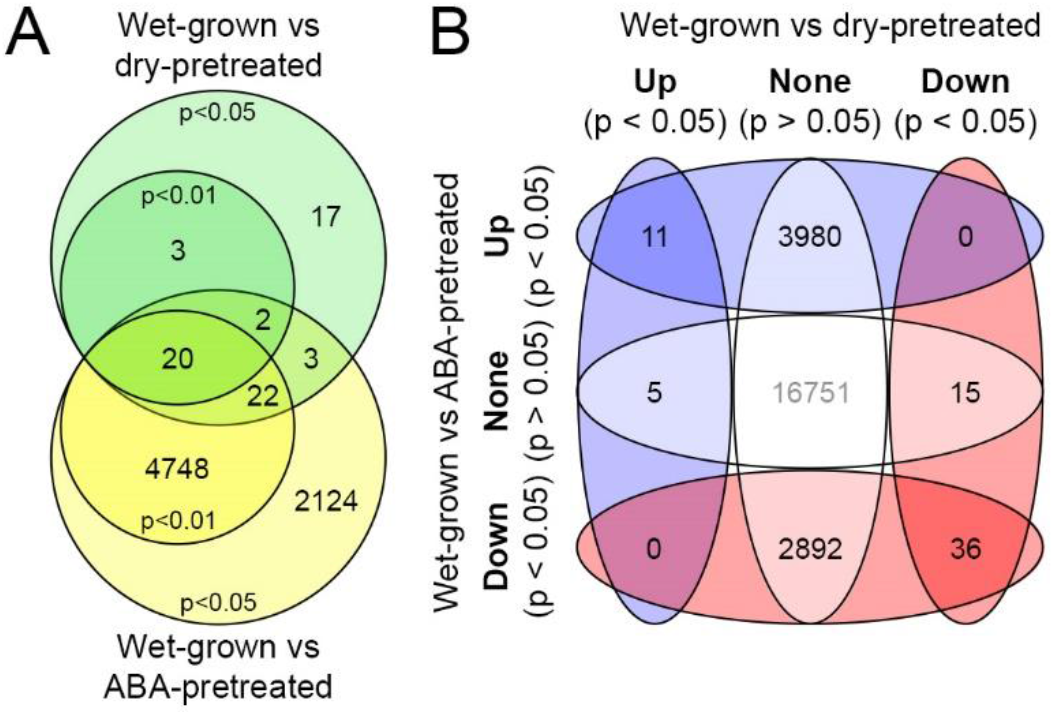
Shared transcriptomic responses in both dry-pretreated and ABA-pretreated *C. richardii* fronds reflect a common underlying sensitisation process. **A.** Venn comparison of genes for which transcript abundance is significantly different between wet-grown and dry-pretreated fronds (green) or wet-grown and ABA-pretreated fronds (yellow) at two levels of stringency (p < 0.01 and 0.01 > p < 0.05, as shown). **B**. Venn comparison between the abundance responses for each transcript identified with a significant change in transcript abundance (p < 0.05) within each pretreatment in (**A**). The central value in grey represents all remaining transcripts within the assembly that do not change abundance in response to either pretreatment.

To assess the likely function of sensitisation signature transcripts, homologs were identified in *Arabidopsis*, similarity to known proteins was determined, and gene ontology (GO) terms were assigned. Eighteen of the 47 sequences have homology to one or more *Arabidopsis* genes, 10 show significant similarity to known proteins in BLAST searches, 12 could be assigned GO terms and 19 have no identifiable similarity to any other sequences (**Dataset S2**). Of those assigned GO terms for ‘biological process’ 46% were associated with responses to biotic or abiotic stimuli, of those assigned for ‘cellular component’ 43% were related to membranes, and of those assigned for ‘molecular function’, 25% were transporters (**Dataset S2**). The majority of transcripts in these categories accumulated to higher levels in wet-grown than in pretreated fronds, indicating down-regulation during sensitisation. Notably, two transcripts are homologous to *Arabidopsis* genes with annotations linked to ABA and stomata, respectively (**SI Appendix**, **Fig. S4**), both encoding plasma membrane ABC transporters that regulate stomatal aperture in *Arabidopsis*. One is homologous to an ABCG subfamily protein (AtPDR12) that transports ABA in stomatal guard cells during closure[^25^] and the second is homologous to an ABCC subfamily protein (AtMRP4) that regulates stomatal aperture either upstream or independently of ABA[^26^]. A third transcript within the sensitisation signature is homologous to AtGRDP1, a negative regulator of ABA responses[^27^]. In both dry- and ABA-pretreated *C. richardii* fronds, the AtPDR12 and AtGRDP1 homologs are both down-regulated whereas the AtMRP4 homolog is up-regulated (**SI Appendix**, **Fig. S4**; **Dataset S2**). Collectively these data suggest that although many transcripts in the sensitisation signature have no known homologs in other species and may have species-specific functions in stomatal regulation, there are recognisable components associated with known stimulus-response signalling mechanisms in flowering plants, including membrane transporters that regulate the inter- and intra-cellular gradients that are crucial for stomatal closure.

### *Homologs of genes required for stomatal closure in* Arabidopsis *are associated with stomatal closure in* C. richardii

To identify genes underlying fern stomatal closure mechanisms, transcript accumulation profiles were generated from fronds with different stomatal sensitivities, before and 60 minutes after exposure to a range of treatments (**Fig. 2*A***; **SI Appendix**, **Fig. S5**). Four treatments were carried out with dry-pretreated (sensitised) fronds: maintenance at high humidity with and without an ABA stimulus (no closure, **Fig. 1*B***), exposure to a low humidity stimulus (closure, **Fig. 1*A***), and simultaneous exposure to low humidity and ABA stimuli (enhanced closure, **Fig. 1*A***). Transcriptomes were also generated from wet-grown (non-sensitised) and ABA-pretreated (sensitised) plants treated with a low humidity stimulus (wet-grown - no closure, **Fig 1*A*;** ABA-pretreated - closure, **Fig 1*C***). For each of the six assays, transcripts were identified that were present at significantly different levels (p <0.01) before and after treatment (**Dataset S3**). Notably, fewer transcripts showed significantly different levels in the assay with ABA-pretreated fronds than in the other assays, possibly reflecting an altered baseline of expression at timepoint 0 as a result of ABA pretreatment, and/or more rapid closure than seen in dry-pretreated stomata (**Fig. 1*C***) precluding the capture of at least some changes underlying the closure mechanism at the sampled 60 minute timepoint. Transcripts specifically associated with stomatal closure were identified by comparing transcript profiles between pairs of assays where stomata closed in one but remained open in the other (**SI Appendix**, **Fig. S5**), using multiple assay-pairs to filter out other experimental or environmental responses. A total of 1858 transcripts that showed a significant change in levels (p < 0.01) specifically upon stomatal closure were identified from four such paired datasets, with the majority being up-regulated on closure (**SI Appendix**, **Fig. S5**). Four of the 1858 transcripts were detected in all four paired datasets, 141 were present in at least three of the datasets and 877 were found in at least two of the datasets (**Fig. 4*A***). Hereafter we refer to the 877 transcripts as stomatal closure-associated transcripts, with the 141 transcripts from within that set that are present in at least three datasets referred to as the stomatal closure signature.

**Figure 4.**
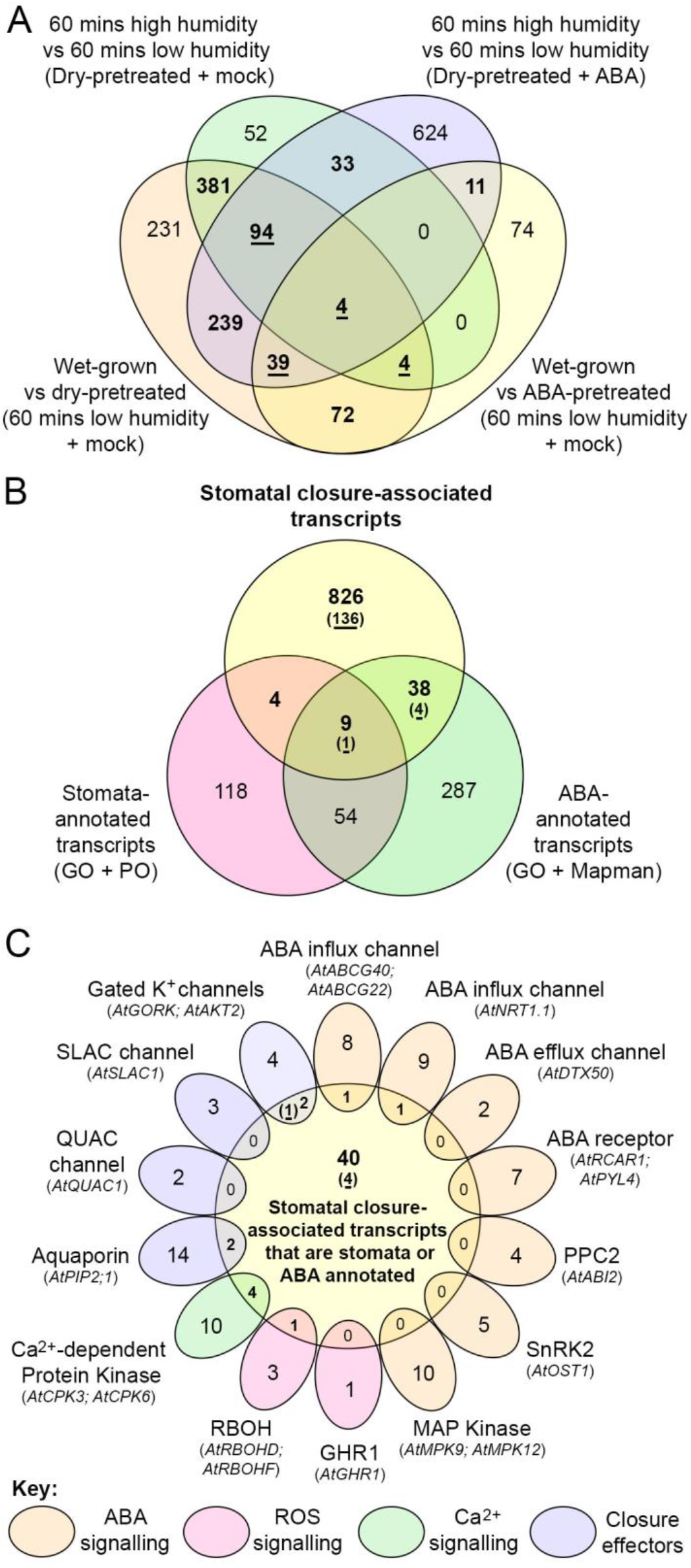
Homologs of genes underlying stomatal closure in *Arabidopsis* are also associated with closure in *C. richardii*. **A.** Venn comparison of transcripts accumulating to significantly different levels (p < 0.01) in association with stomatal closure between four different closure datasets (as shown). Within each dataset, transcripts specific to stomatal closure were identified by comparison with similar experimental conditions in which stomata did not close (**SI Appendix**, **Fig. S6**). A total of 877 transcripts are associated with closure in at least two datasets (bold), 141 of which are present in at least three datasets and are thus referred to hereafter as closure signature transcripts (bold, underlined). **B.** Venn comparison between the 877 closure-associated transcripts (labelled in bold in **A**), and all assembled transcripts with homology to *Arabidopsis* genes with ABA or stomata-related functions based on annotation in Gene Ontology, Plant Ontology or Mapman databases (as shown). Underlined numbers given in brackets denote the 141 closure signature transcripts (underlined in **A**) within each category, where present. A total of 51 closure-associated transcripts overlapped with homologs of stomata or ABA-annotated genes. **C.** Venn comparison between the 51 closure-associated transcripts annotated as ABA and/or stomata-related in (**B**) and all transcripts within the assembly homologous to regulators of stomatal closure in *Arabidopsis*. Components of different regulatory pathways (ABA signalling, Reactive Oxygen Species [ROS] and calcium signalling) and final effectors are colour-coded, as indicated. The *Arabidopsis* genes used to determine homology for each gene family/functional category are listed in brackets. Underlined numbers given in brackets denote the 5 members of the signature subset identified in (**B**) within each category, where present.

To determine whether any of the closure associated transcripts are homologous to genes involved in *Arabidopsis* stomatal closure networks, stomatal and ABA annotations were mapped onto the whole transcriptome assembly (see Materials and Methods). Fifty-one of the 877 transcripts had homology to *Arabidopsis* genes with stomatal or ABA annotations, of which five were closure signature transcripts (**Fig. 4*B***). To further assess possible gene function, the 51 transcripts were screened for homology to 14 *Arabidopsis* genes with proven roles in stomatal closure[^2,28^], all 14 of which had identifiable homologs within the whole transcriptome assembly (**Dataset S4**). These comparisons revealed that one of the five annotated signature transcripts was homologous to the SHAKER family of gated potassium ion channels that function in the guard cell membrane[^29^] (**Fig. 4*C***). Homologs of aquaporin, CPK calcium-dependent protein kinase, respiratory burst oxidase homolog (RBOH) and annexin gene families were present within the remaining group of 46 annotated closure-associated transcripts (**Fig. 4*C***; **Dataset S3**). Although no canonical components of the ABA signal transduction pathway were identified, two families of ABA influx channels involved in stomatal closure were also represented (**Fig. 4*C***). Collectively, these data show that homologs of genes in many of the signalling pathways that mediate stomatal closure in *Arabidopsis* are associated with stomatal closure in *C. richardii*, and thus suggest that closure mechanisms are at least partially conserved in ferns and angiosperms.

### *Changes in transcript abundance during stomatal closure and sensitisation in* C. richardii *are largely ABA-independent*

To further dissect ABA-related mechanisms underlying stomatal closure in *C. richardii*, we exploited the fact that stomata in dry-pretreated plants only close in response to ABA when the ABA treatment is carried out in conditions of low humidity (**Fig. 1*A,B***). ABA-related changes in transcript abundance that are specific to stomatal closure should thus be observed following ABA treatment under low humidity but not following treatment under high humidity. Transcript levels that changed in response to ABA under low and high humidity conditions were identified by comparing transcriptomes of fronds that were exposed to either a mock or ABA treatment solution (**SI Appendix**, **Fig. S6**). The abundance of 578 and 607 transcripts was significantly altered (p < 0.01) in response to ABA under low and high humidity conditions, respectively. Levels of two of the 141 closure signature transcripts changed after ABA treatment under high humidity (representing an unknown protein and a UDP-glucose 6-dehydrogenase, **Dataset S5**), but none changed after treatment under low humidity (**Fig. 5*A***). Abundance of a single closure-associated transcript (representing a polyphenol oxidase) was influenced by ABA under both humidity conditions but levels changed in opposite directions in the two conditions (**SI Appendix**, **Fig. S6**; **Dataset S5**). Levels of just 13 closure associated transcripts changed in response to ABA treatment specifically under low humidity (**SI Appendix**, **Fig. S6**; **Dataset S5**). These results suggest that ABA-induced closure of fern stomata (**Fig. 1**) is mediated primarily via mechanisms operating at translational or post-translational levels.

**Figure 5.**
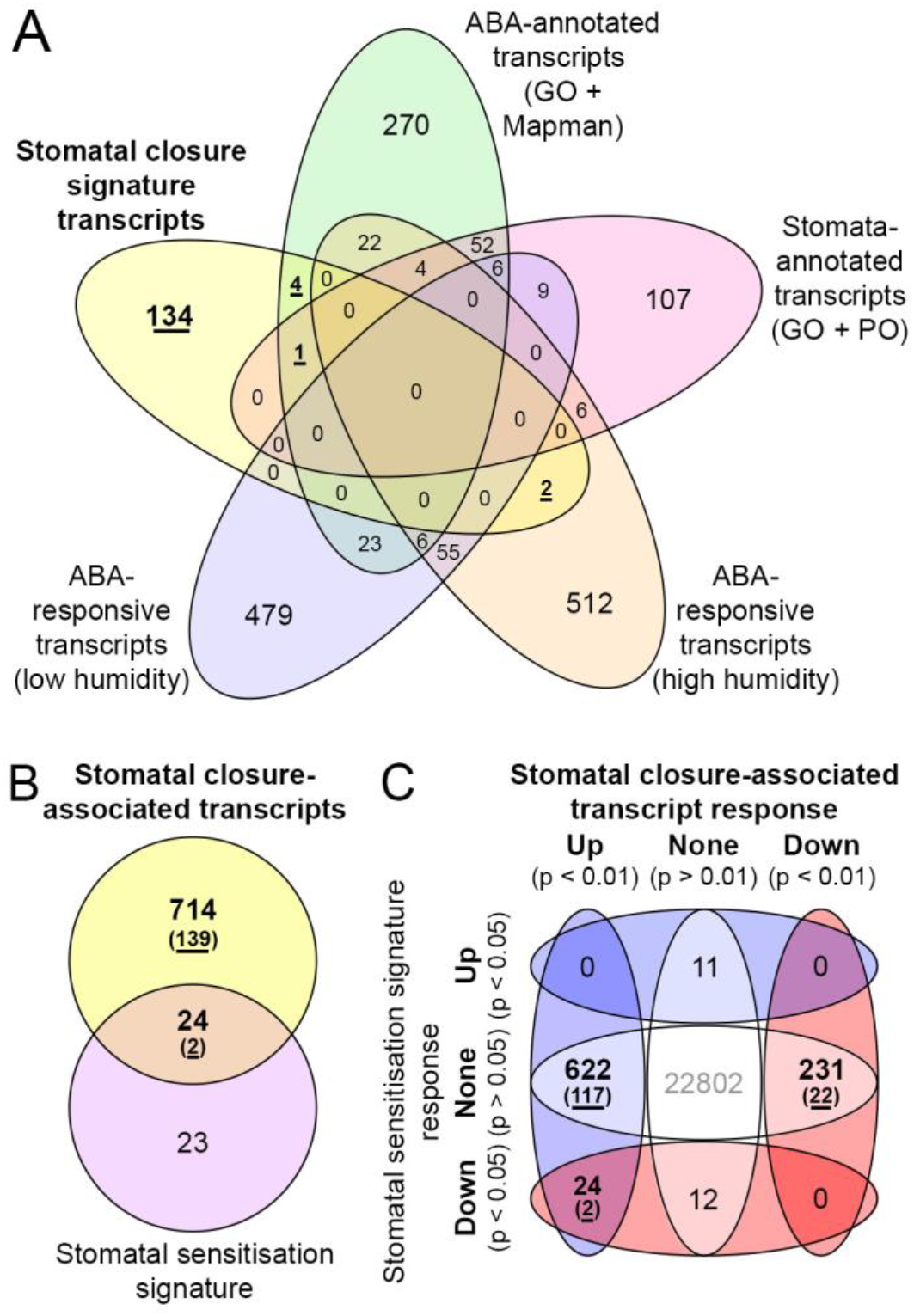
Transcriptome profiles during stomatal closure and sensitisation suggest shared but opposing regulatory mechanisms that are largely ABA-independent. **A**. Venn comparison between stomatal closure signature transcripts (bold, underlined, see **Fig. 4*A***), all transcripts with annotations relating to ABA and stomata functions, and all transcripts found to significantly change abundance (p < 0.01) in response to ABA treatment under low or high humidity. Transcript levels that changed specifically in response to ABA were identified by comparing abundance at 0 and 60 minutes between mock and ABA treatments (**SI Appendix**, **Fig. S6**). No overlap between the stomatal closure signature and ABA-responsive transcripts was found. **B.** Venn comparison between transcripts identified as stomatal closure-associated transcripts (bold) and the stomatal sensitisation signature (see **Fig. 3*A***). Underlined numbers given in brackets denote closure signature transcripts from within the closure-associated transcript dataset. **C.** Venn comparison between the directional change in level of stomatal closure-associated (bold) and sensitisation signature transcripts during stomatal closure and stomatal sensitisation, respectively. Underlined numbers given in brackets denote closure signature transcripts within the closure-associated transcript dataset. Transcripts common to both sensitisation and closure datasets are all down-regulated in sensitised fronds relative to wet-grown controls and are all up-regulated during closure.

### *Transcriptome profiles during stomatal sensitisation and closure in* C. richardii *suggest that sensitisation is associated with the down-regulation of closure-associated genes*

To determine whether stomatal sensitisation and closure mechanisms share any common features, the identities of sensitisation signature transcripts were compared to those of closure associated transcripts. Twenty four of the 47 sensitisation signature transcripts were present in the closure associated transcript dataset, with two also being present in the much smaller closure signature dataset (**Fig. 5*B***). Fourteen of these 24 common transcripts have similarity to known proteins (**Dataset S3**), including homologs of the ABCG transporter AtPDR12 (**Fig. 4*C***), AtGRDP1, a peroxidase superfamily protein, and two calcium-binding EF-hand family proteins. Notably, all 24 transcripts decreased in abundance during sensitisation and increased in abundance during closure (**Fig. 5*C***), returning to a similar level as seen in untreated wet-grown plants (p > 0.05; **SI Appendix**, **Fig. S7**; **Dataset S3**). The fact that more than 50% of the sensitisation signature transcripts are also stomatal closure associated transcripts suggests that stomatal sensitisation acts by priming the closure mechanism.

## Discussion

Stomata facilitated the adaptation of plants to land and with the exception of liverworts, all extant land plant lineages rely on stomatal function to modulate the exchange of air and water with the surrounding environment. Despite the likely monophyletic origin of stomata and their conserved function across these extant groups, the evolution of their regulatory networks is disputed, with discussions focused on whether the ABA-dependent active stomatal closure mechanisms found in flowering plants[^2,30^] are conserved in non-flowering plant lineages. We show that stomatal closure in the model fern *C. richardii* is active, but is conditional on sensitisation by prior exposure to either low humidity or ABA. To identify the mechanisms underlying these processes, we utilised RNA-seq to compare transcriptomes of whole fronds that were exposed to different conditions under which stomata did or did not close, generating a high-quality *de novo* assembly from which we could discriminate signatures associated with stomatal responses. These show that both stomatal sensitisation and closure in *C. richardii* include changes in the expression of genes required for active stomatal closure in the flowering plant *Arabidopsis*, suggesting that guard cell signalling mechanisms are conserved. Active stomatal closure responses are thus likely to have evolved in the last common ancestor of land plants.

### Conserved active stomatal closure mechanisms in ferns and flowering plants

The requirement for sensitisation to reveal active stomatal closure responses, shown here in *C. richardii* and previously in some other fern species[^17^], might reconcile past reports of non-active stomatal responses in basal land plant lineages[^10,11,12,13,14^] but the extent to which conditional stomatal sensitivity is present across the fern lineage remains unclear. Humidity-conditional stomatal responses to ABA have been reported in two other leptosporangiate fern families[^17^] but eusporangiate ferns have not yet been investigated. Fern stomatal responses to ABA without prior sensitisation have also been reported[^9,20^] but in contrast one fern species did not show closure or ABA responses despite dry pretreatment[^17^]. Widespread utilisation of active responses in ferns is indirectly supported, however, by a recent study showing that stomata in species from across the fern lineage open in response to blue light[^31^], as seen in flowering plants[^32^]. Our findings thus contribute to an emerging consensus that active regulation of stomatal responses is present across the fern lineage.

ABA induction of stomatal closure in *Arabidopsis* involves activity of the canonical RCAR/PRY/PYL - PP2C ABA signal transduction complex[^30,33,34,35^], two well-characterised ABA responsive proteins (Snf1-Related Kinase subfamily 2 (SnRK2)[^36^] and S-type anion channels (SLAC)[^37,38^]) and a number of downstream genes including those encoding ABA influx channels, calcium and ROS signalling components, and K^+^ ion channels[^2^]. Although *C. richardii* homologs of genes in many of the downstream families were identified in the stomatal closure-associated dataset generated here, RCAR/PRY/PYL, SLAC and SnRK2 homologs showed no change in transcript levels upon stomatal closure, despite ABA treatment causing more rapid closure responses. Given that stomatal SLAC channel activity is activated post-transcriptionally by ABA in *Arabidopsis*, via SnRK2-mediated protein phosphorylation[^39,40,41^], our transcriptome data do not discount a similar post-transcriptional mechanism operating during stomatal closure in *C. richardii*. Furthermore, although a previous study of individual SnRK2 and SLAC gene function in *C. richardii* failed to find evidence for such a role[^21^], our transcriptome assembly has identified additional SnRK2 and SLAC homologs (**Dataset S4**), raising the possibility of functional redundancy. With recent evidence suggesting that stomata of leptosporangiate ferns have acquired greater responsiveness to blue light through duplicated CRY receptors[^31^], the presence of multiple SnRK2 and SLAC gene copies even raises the possibility of a more complex mechanism. Although conservation of SnRK2-SLAC interactions across land plants has previously been supported by mutant and *in vitro* functional analysis in the moss *Physcomitrella patens*[^7,8^], further *in vivo* testing is clearly required to resolve whether the SnRK2-SLAC pathway functions in *C. richardii* stomatal closure. Regardless, our results suggest that downstream components of active stomatal closure mechanisms are conserved in flowering plants and ferns, either fully or partially, and thus that a similar molecular framework was operating in guard cells of the last common ancestor of the two lineages.

### Stomatal sensitisation in C. richardii shares features with stomatal acclimation in flowering plants

Both dry and ABA pretreatment are sufficient to condition stomatal sensitivity in *C. richardii*, with both treatments inducing similar genome-wide changes in transcript abundance that enable an active closure response on subsequent exposure to low humidity. The stomata of flowering plants can also be sensitised or de-sensitised to closure-inducing signals, through the process of acclimation. For example, newly-developed *Arabidopsis* stomata become increasingly sensitive to closure-promoting environmental stimuli during leaf maturation in an ABA-dependent manner[^42^] whereas species transferred to high humidity conditions show impaired stomatal closure when subsequently exposed to lower humidity, despite accumulation of endogenous or exogenous ABA[^43,44^]. In flowering plants, the mechanism underlying altered guard cell sensitivity to ABA is thought to involve altered abundance of the ABA receptors PYL/PYR/RCAR[^30^]. Although there was no evidence of increased ABA receptor abundance in sensitised *C. richardii* fronds, the possibility that the whole-frond dataset was not sufficiently sensitive to detect all guard cell-specific changes cannot be excluded. However, reduced abundance of an AtGRDP1 homolog[^27^] was detected in sensitised fronds. Given that AtGRDP1 negatively regulates the expression of the ABA response regulators *ABI3*[^45^] and *ABI3*[^46^], and that loss of function leads to ABA hypersensitivity[^27^] it is thus possible that stomatal sensitisation in *C. richardii* results from increased sensitivity to ABA. As such, conditional stomatal sensitivity in ferns and stomatal acclimation in flowering plants may both act by altering the threshold at which subsequent inductive signals, including ABA, can trigger stomatal closure.

Consistent with the idea that stomatal sensitisation lowers the threshold for induction of stomatal closure, the abundance of 24 closure-associated transcripts is reduced during sensitisation, with all 24 returning to the same levels seen prior to sensitisation upon subsequent induction of closure. This set of 24 includes all transcripts in the sensitisation signature that have calcium-binding and ROS-related annotations, plus homologs of AtGRDP1 and AtPDR12, which is an ABCG transporter that increases guard cell uptake of ABA[^25^]. On the basis of these observations, we speculate that sensitisation results when Ca^2+^, ROS and/or ABA signalling cascades within the guard cell are potentiated to reduce the threshold required for a stimulus to trigger closure. Alongside this down-regulation of closure-associated genes, it is notable that a putative homolog of AtMRP4 is conversely up-regulated during sensitisation. AtMRP4 is a guard cell-expressed ABCC transporter that negatively regulates stomatal opening in an ABA-independent manner[^26^]. Collectively, our results support a scenario whereby fern stomata are sensitised by altering the intracellular physiological status of guard cells, enhancing the ability to respond to canonical closure signals, and attenuating competing opening responses, using mechanisms conserved with flowering plants.

## Materials and Methods

### Plant material and growth conditions

All experiments were performed using *Ceratopteris richardii* wild-type laboratory strain Hn-n[^47^]. Gametophyte and sporophyte phases were grown under sterile culture on ‘C-fern’ solid growth media[^48^] (1% agar, pH6.0) under 16hrs light/ 8 hrs dark photoperiod at a fluence of 150μmol m^−2^ s^−2^ and constant 28°C temperature. Synchronised populations of sporophytes were generated for experiments by free mating of gametophytes as previously described[^49^]. Individual sporophytes were subsequently transferred to GA-7 Magenta boxes (Merck Life Science UK Ltd., Dorset, UK) containing 100ml C-fern agar media when two fronds were visible (10-14 days old). Humidity was maintained by the application of 1.5-2ml sterile water at the time of transplant, and subsequently when no surface water was visible (approximately every 14 days), with the time each box was open minimised to maintain high humidity.

Sporophytes were grown to 46-50 days old before sampling fronds for both stomatal response assays and RNA-seq. Plants were pretreated with either periodic low humidity or exogenous ABA application for a period of three weeks after transplant, then grown without further pretreatment for two weeks. During pretreatment, humidity conditions within magenta boxes were measured across three replicates using a TinyTag 2Plus (Gemini Data Loggers, Chichester, UK). A constant high humidity of 94.9±0.36% (± S.E.) was recorded within unopened Magenta boxes. ‘Dry’ pretreatment was applied on alternating days three times a week by opening Magenta boxes to ambient humidity (48.3 ± 0.69%) under sterile conditions for 10 minutes, during which time humidity within the boxes fell to 78.8±2.11%. After boxes were closed internal humidity returned to ≥ 90% in 3.3±0.33 minutes. Plants under stable high humidity (‘wet-grown’) pretreatment were incubated alongside dry-pretreated magenta boxes during their exposure but were kept closed. ABA and mock pretreatments were delivered to plants inside magenta boxes as 100μM exogenous ABA in buffered solvent (10mM MES/KOH pH6.2, 0.5% Tween20 (v/v)) or a corresponding mock solution (0.1% EtOH in buffered solvent) respectively, applied dropwise twice a week with the time each box was open minimised to maintain high humidity. Opening of magenta boxes for exogenous treatment or water application caused a transient fall in recorded humidity to a minimum of 86.4±0.93%.

### Stomatal response assay

The responses of stomata from humidity- and ABA-pretreated plants were tested in separate assays. Plants undergoing stomata response assays were treated with 100uM ABA or mock solutions (formulation as pretreatment) at the start of the assay (0 minutes), applied by foliar spray in a design orthogonal to humidity or ABA pretreatments such that each pretreatment+treatment combination was represented by three plants per assay. Plants were either immediately exposed to low humidity by removal from Magenta boxes into ambient air (48.3±0.69%), or in control experiments maintained at high humidity by immediately resealing Magenta boxes after ABA/mock treatment. At three timepoints (0, 60 and 120 minutes from the assay start) a single frond was sampled from one plant for each pretreatment+treatment combination and photographed to record stomata morphology, sampling an independent plant at each timepoint within each pretreatment+treatment combination, selecting fronds at shoot position 7-9, bearing 3 lobes (**SI Appendix**, **Fig. S1**). Stomata morphology was recorded by imaging was under brightfield microscopy at 10x magnification using an Olympus BX51 microscope (Olympus Corporation, Hamburg, Germany) with attached Qimaging MP3.3-RTV-R-CLR-10-C MicroPublisher camera and QImage Pro software package (Teledyne Photometrics, Tuscon, USA). Photography at each timepoint was completed within 15 minutes, and pretreatment+treatment combinations were imaged in the same order between timepoints. Three separate regions were photographed on each frond (one within each lobe), and 20 stomata within each region were selected systematically for measurement, totally 60 stomata per plant per timepoint. Stomatal measurements were made from scaled images using the FIJI software package[^50^]. Assays were repeated three times for each combination of pretreatments and low or high humidity stimulus. Whole-plant photography was performed using a Cybershot DSC-HX7V digital camera (Sony, Tokyo, Japan). During figure preparation images were adjusted for brightness and contrast using Photoshop 2019 (Adobe, San Jose, USA).

### RNA extraction and sequencing

The sampling of frond tissue for RNA-seq was performed by snap-freezing single fronds in liquid nitrogen from specific pretreatment+treatment combinations at 0 or 60 minutes (**Fig. 2*A***). Three biological replicates were harvested for each pretreatment+treatment combination, taken from three independently-performed assays. RNA was extracted using the RNeasy Plant mini kit with on-column DNase treatment (QIAgen, Hilden, Germany). RNA quality was validated by Agilent 2100 Bioanalyser RNA 6000 Pico assay (Agilent, Santa Clara, USA). Libraries were prepared from single frond samples using 1μg starting RNA using the TruSeq RNA library prep kit V2 (Illumina Systems, San Diego, USA) and quantified by KAPA library quantification (Roche, Basel, Switzerland) on a CFX96 qPCR machine (Bio-Rad, Hercules, USA). Paired-end sequencing was performed by HiSeq2000 sequencing system using a NextSeq 500/550 High Output kit (150 cycles) (Illumina Systems, San Diego, USA).

### Transcriptome assembly and analysis

Paired end reads were processed using rCorrector[^51^] with default parameters to correct likely erroneous k-mers. The Harvard FAS Informatics “FilterUncorrectabledPEfastq” script (https://github.com/harvardinformatics/TranscriptomeAssemblyTools, accessed 13 January 2020) was used to discard read pairs for which one of the reads was deemed unfixable. TrimGalore (https://www.bioinformatics.babraham.ac.uk/projects/trim_galore) was used with default settings to trim adapter sequences and low-quality bases identified using the tools Cutadapt[^52^] and FastQC[^53^]. Reads were mapped to the SILVA rRNA SSUParc and LSUParc databases, release 132[^54^] using Bowtie 2[^55^] with option “--very-sensitive-local” and those pairs for which neither read was mapped to the rRNA database were retained. Analysis using FastQC showed that the retained reads had per base sequence content deviations greater the 5% from the mean for the first 7 and last 3 bases of the reads and so these were clipped using TrimGalore. The transcriptome was assembled using Trinity[^56^] with default settings. Transcript abundance estimation was performed using Salmon[^57^] within the Trinity pipeline. Differential expression testing was performed using DESeq2[^58^] (options “-min_reps_min_cpm 2,1”) also with the Trinity pipeline. All differential expression comparisons between sample pairs are given in **Datasets S2**, **S3** and **S5**. Venn diagrams were generated using the InteractiVenn web tool[^59^].

A proteome was constructed by analysing the transcriptome using Transdecoder (http://transdecoder.sf.net) “LongOrfs” and “Predict” with the Trinity gene-to-transcript map supplied as input and with default parameters. Amino acid sequences were additionally identified for sequences that were expressed above the minimum threshold for the differential expression analysis (> 1 CPM in at least two biological replicates) but which had not yet had an amino acid sequence identified. This was achieved using “TransDecoder.LongOrfs” with option “-m 10” to set a minimum length of 10 amino acids. Those transcript sequences that did not return an open reading frame using TransDecoder were subject to open reading frame prediction using GeneMark-ES[^60^].

To aid in the annotation of the differentially expressed transcripts, Longest transcript variant proteomes for the 12 species: *Arabidopsis thaliana* (TAIR10)[^61^], *Amborella trichopoda* (v1.0)[^62^], *Glycine max* (Wm82.a4.v1)[^63^], *Manihot esculenta* (v7.1)[^64^], *Micromonas pusilla* CCMP1545 (v3.0)[^65^], *Ostreococcus lucimarinus* (v2.0)[^66^], *Oryza sativa* (v7.0)[^67^], *Physcomitrella patens* (v3.3)[^68^], *Solanum lycopersicum* (ITAG3.2)[^69^], *Selaginella moellendorffii* (v1.0)[^70^], *Spirodela polyrhiza* (v2)[^71^] and *Volvox carteri* (v2.1)[^72^] were downloaded from Phytozome[^73^]. OrthoFinder 2.3.11[^74^] was used with default settings to assign the genes to orthogroups and to identifying orthologs for the *C. richardii* proteome. GO[^75,76^], Plant Ontology[^77^] and MapMan[^78^] annotations for *Arabidopsis* were downloaded from The Arabidopsis Information Resource (TAIR)[^61^]. Each orthogroup inherited the annotations of the *Arabidopsis* genes that it contained. The same reference proteomes were used to assess completeness of the predicted *C. richardii* proteome using BUSCO analysis[^79^].

To identify homologs of the annotated protein coding genes a series of searches were carried out, terminating at the first success for each gene. These searches were for: an ortholog in *Arabidopsis* identified by OrthoFinder, an *Arabidopsis* homolog from within the same OrthoFinder orthogroup, any significant DIAMOND[^80^] hit to an *Arabidopsis* homolog (e < 10^−3^), and significant hit for a homolog within the 12 plant species above, any significant hit within the BLAST nr database[^81^].

### Data availability

The sequencing reads obtained from each library have been deposited in the NCBI Sequence Read Archive (SAMN16306556) and the assembled transcriptome has been deposited in the NCBI Transcriptome Shotgun Assembly Sequence Database under Bioproject PRJNA666635. The predicted proteome is available on figshare at the link:https://doi.org/10.6084/m9.figshare.13807967.

## Author contributions

ARGP performed stomatal response assays, RNA extraction and next-generation sequencing library preparation. DME performed transcriptome assembly and bioinformatic analysis. ARGP, AMH, JAL and SK designed experiments. All authors contributed to preparation of the manuscript.

## Acknowledgements

This work was supported by an ERC Advanced Investigator Grant (EDIP) to JAL, by a grant from the Gatsby Charitable Foundation to ARGP and AMH, and by Royal Society University Research Fellowships to ARGP and SK. SK and DE were supported by the European Union’s Horizon 2020 research and innovation programme under grant agreement number 637765. AMH would like to acknowledge support from the Leverhulme Trust, RPG-2019-004 and the New Phytologist Foundation We are grateful to Julie Bull and Ester Rabbinowitsch for support with plant watering, Prof. Julian Hibberd for providing growth space for RNA-seq experiments and to Pawel Baxter for performing NGS sequencing runs.

## Supplemental Information

### SI Appendix

**Fig. S1.** Effect of pretreatments on stomata, frond and whole plant morphology in *C. richardii*

**Fig. S2.** Quantification of stomatal dimensions in response to different pretreatment and treatment combinations.

**Fig. S3.** Phenotype of *C. richardii* plants after exposure to low humidity and ABA stimuli following different pretreatments.

**Fig. S4.** Two transcripts associated with stomatal sensitisation in *C. richardii* are annotated with ABA- or stomata-related functions.

**Fig. S5.** Transcriptome changes associated with stomatal closure in *C. richardii*.

**Fig. S6.** Humidity-dependent ABA regulation of stomatal closure-associated and stomatal sensitisation signature transcripts.

**Fig. S7.** Closure-associated transcripts down-regulated by sensitisation return to their original levels during or after stomatal closure.

**Dataset S1.***C. richardii* stomatal response measurements.

**Dataset S2.***C. richardii* stomatal sensitisation signature transcripts.

**Dataset S3.***C. richardii* stomatal closure-associated transcripts.

**Dataset S4.***C. richardii* transcripts homologous to *Arabidopsis* stomata closure genes.

**Dataset S5.***C. richardii* transcripts responsive to ABA under either low or high humidity.

## Notes

https://doi.org/10.6084/m9.figshare.13807967

### Competing Interest Statement

The authors have declared no competing interest.

