## Supplemental Figures S1-S7 for "Conditional Stomatal Closure in a Fern Shares Molecular Features with Flowering Plant Active Stomatal Responses"

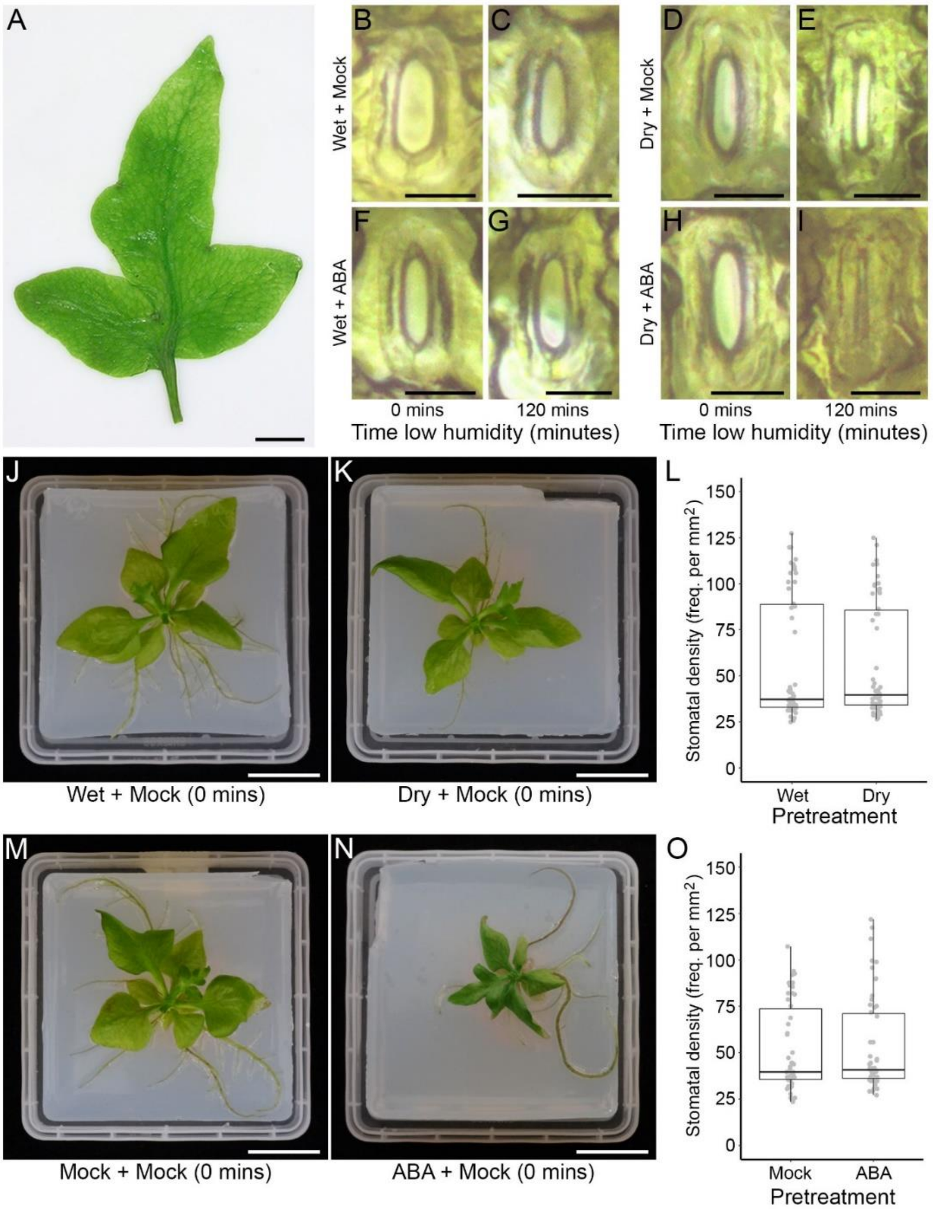


**Fig. S1.** Effect of pretreatments on stomata, frond and whole plant morphology in *C. richardii*

**A.** Morphology of the *C. richardii* frond (position 9-11 on shoot) used for stomatal response assays. Scale bar = 10 mm.

**B-I.** Stomatal morphology on fronds from wet-grown (Wet; **B-C, F-G**) or dry-pretreated (Dry; **D-E, H-I**) plants that were treated at 0 minutes with either mock solution (**B-E**) or 100µM exogenous ABA (**F-I**), and then exposed to a low humidity stimulus for 120 minutes. Scale bars = 25µm.

**J-L.** Effects of pre-treatments on whole plant morphology. **J,K**, morphology of 50 day-old wet-grown (**J**) or dry-pretreated (**K**) plants at 0 minutes exposure to a low humidity stimulus. **L**, stomatal density on fronds from wet-grown or dry-pretreated plants used for stomatal response assays in **Fig. 1*A,B***.

**M-O**. Effects of ABA pre-treatments on whole plant morphology. **M,N**, morphology of 50 day-old wet-grown plants pretreated with mock solution (**M**) or 100µM ABA (**N**) at 0 minutes exposure to a low humidity stimulus. **O**, stomatal density on fronds from mock- or ABA-pretreated plants used for stomatal response assays in **Fig. 1*C,D***.

*N* = 18 independent plants, 3 density measurements taken across a single frond lamina per plant. Boxplots represent the 25$^{th}$-75% percentile (box), the median value (mid-line) and up to 1.5x the interquartile range (whiskers) (**L, O**). Scale bars = 20mm (**J,K,M,N**).

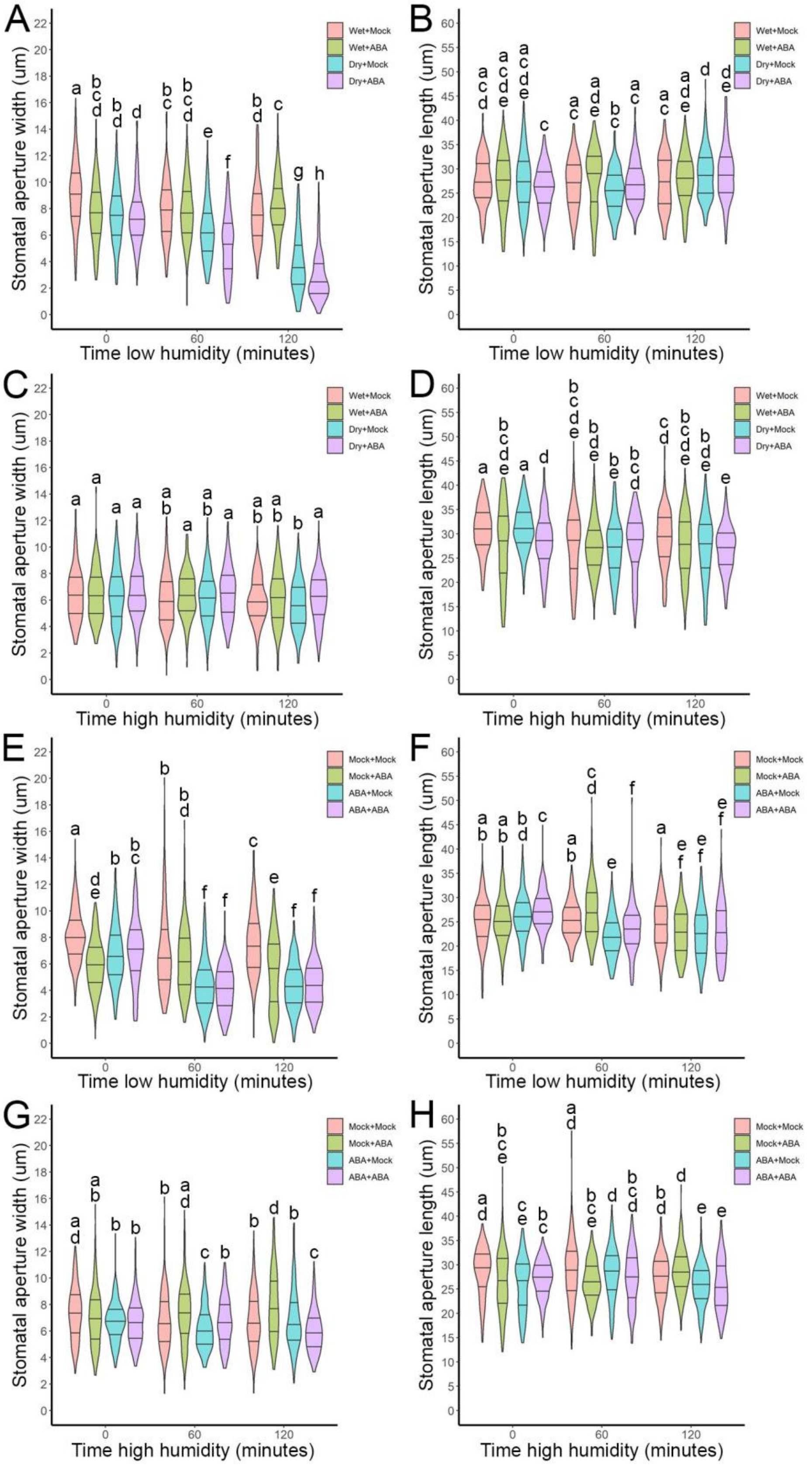


**Fig. S2.** Quantification of stomatal dimensions in response to different pretreatment and treatment combinations.

**A-H**. Stomatal dimensions in fronds from wet-grown or dry-pretreated plants (**A-D**) and wet-grown plants pretreated with periodic exogenous application of 100uM ABA (‘ABA’) or corresponding mock solution (‘Mock’) (**E-H**). Fronds were either exposed to a prolonged low humidity stimulus (**A,B,E,F**) or maintained in high humidity conditions (**C,D,G,H**). Dimensions shown are stomatal width (distance between the midpoint of the two guard cells) (**A,C,E,G**) and length (distance between the points where the guard cells attach) (**B,D,F,H**). Either mock solution (‘+Mock’) or 100uM exogenous ABA (+ABA) was applied in an orthogonal design at the start of the assay.

*N* = 180 stomata per pretreatment + treatment combination, measured from three independent plants per timepoint. Violin plots represent data distribution density, with thickness corresponding to frequency of datapoints at that value. Median value and upper and lower quartiles are represented by the middle, upper and lower horizontal lines, respectively. Pairwise comparisons were performed using two-tailed Mann-Whitney tests. Where necessary to meet the assumptions of this test, comparisons were performed on square-root (**E**,**F**,**G**) or $\log_{10}$-transformed datasets (**H**). Letters denote statistically significant differences ($p < 0.05$) within each plot. Pretreatment + treatment combinations with the same letter are not significantly different from each other.

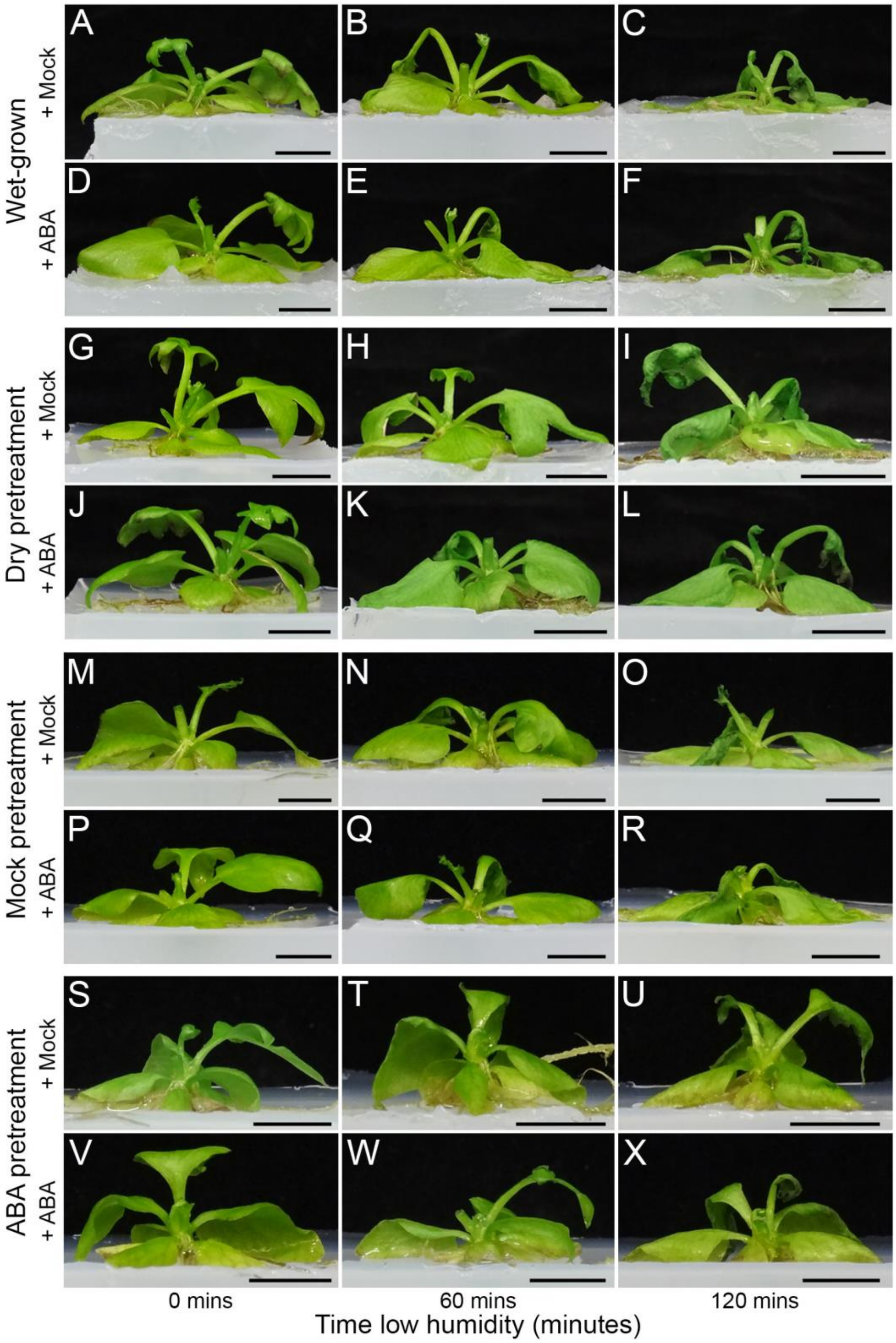


**Fig. S3.** Phenotype of *C. richardii* plants after exposure to low humidity and ABA stimuli following different pretreatments.

Whole-plant morphologies of wet-grown (**A-F**), dry-pretreated (**G-L**), mock-pretreated (**M-R**) and ABA-pretreated (**S-X**) plants, 0, 60 and 120 minutes after transfer to a low humidity environment from an initial stable wet-growth environment. Within each pretreatment group, plants were treated at 0 minutes with mock solution (**A-C**, **G-I**, **M-O**, **S-U**) or 100uM ABA (**D-F**, **J-L**, **P-R**, **V-X**) by foliar spray. Scale bars = 10mm.

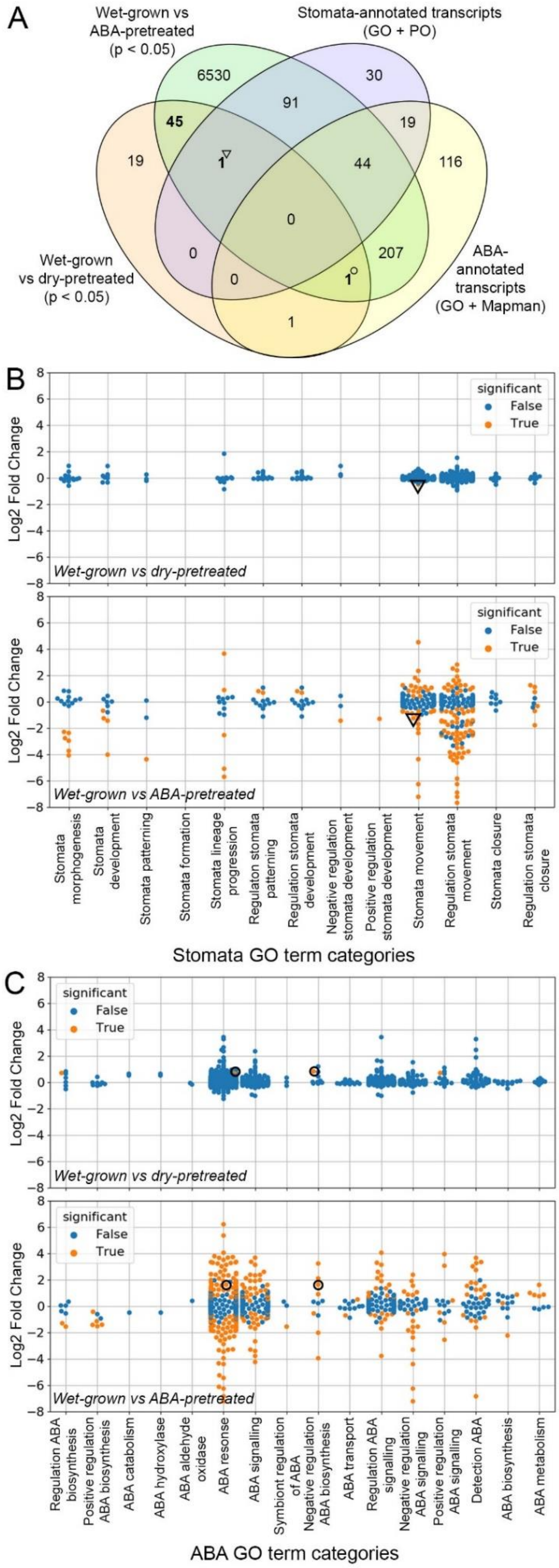


**Fig. S4.** Two transcripts associated with stomatal sensitisation in *C. richardii* are annotated with ABA- or stomata-related functions.

**A.** Venn comparison between transcripts with altered abundance ($p < 0.05$) in response to dry pretreatment or ABA pretreatment (as shown) and all assembled transcripts with homology to *Arabidopsis* genes with ABA or stomata-related functional annotations from Gene Ontology (GO), Mapman and Plant Ontology (PO) databases (as shown). Two single transcripts in the stomatal sensitisation signature dataset (highlighted in bold) overlap separately with ABA (TRINITY_DN94_c0_g1; AtPDR12 homolog) and stomata-annotated transcripts (TRINITY_DN8731_c0_g1; AtMRP4 homolog), marked with an open circle and open triangle, respectively.

**B,C.** Comparison of abundance level changes after dry pretreatment or ABA pretreatment (as shown) for all *C. richardii* transcripts with GO term annotations relating to stomata (**B**) or ABA (**C**). The level of each transcript is expressed as a fold-change of the wet-grown control relative to the pretreatment applied, on a log2-transformed scale. Colour coding indicates whether the transcript level change is significant (orange, $p < 0.05$) or not significant (blue, $p \geq 0.05$). The datapoints marked with an open circle and open triangle represent the two transcripts identified in (**A**) and show similar responses between the two pretreatments. Transcript TRINITY_DN94_c0_g1 (open circle) is annotated with more than one ABA GO term category.

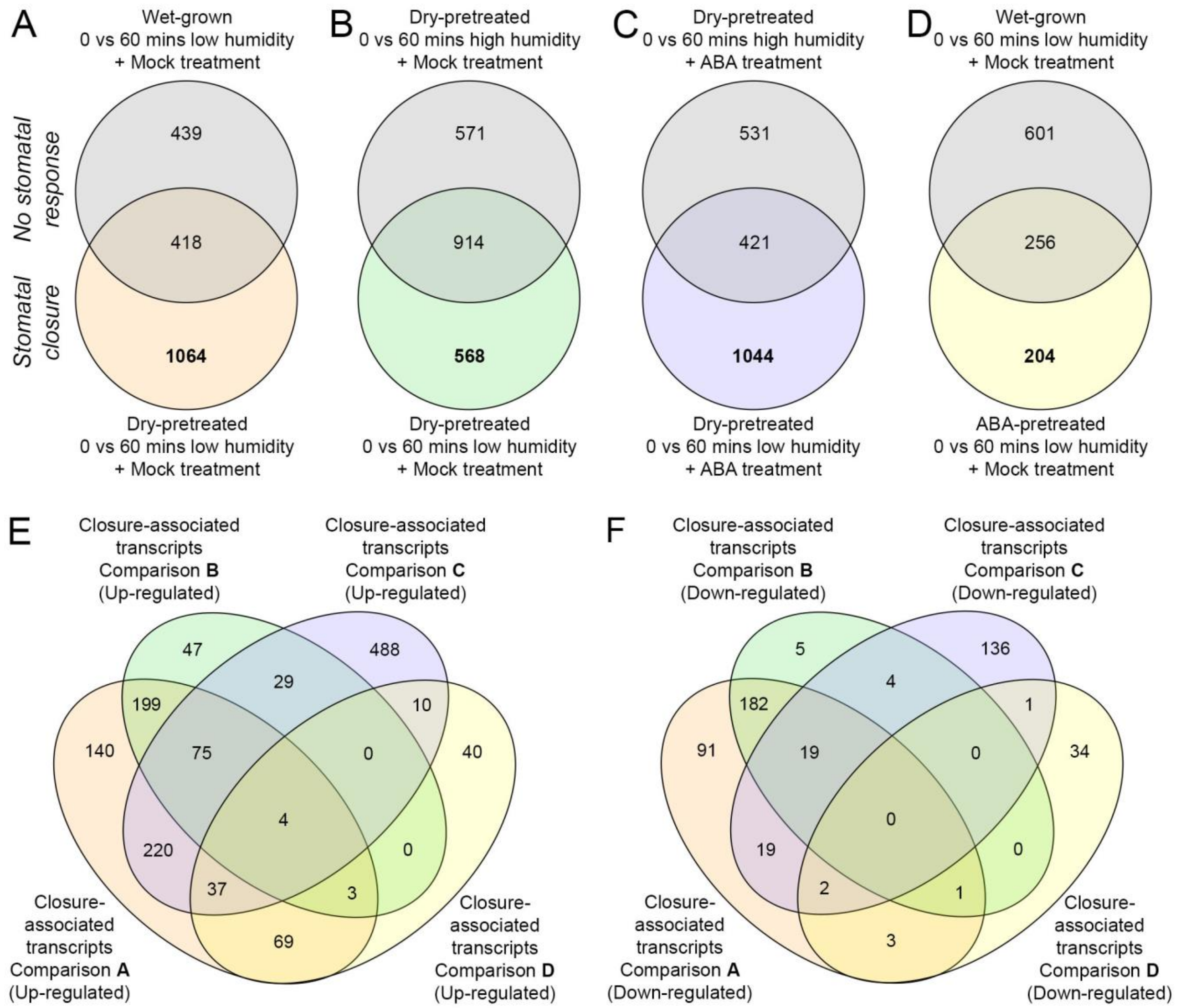


**Fig. S5.** Transcriptome changes associated with stomatal closure in *C. richardii*.

**A-D.** Venn comparisons of transcripts that accumulated to significantly different levels ($p < 0.01$) in stomatal response assays. For each assay, transcripts with significantly different abundance levels ($p < 0.01$) in 0 and 60 minute samples were identified. This revealed significant differences for: 857 transcripts in wet-grown, low humidity treated samples; 1485 in dry-pretreated, high humidity treated; 952 in dry-pretreated, high humidity & ABA treated; 1482 in dry-pretreated, low humidity treated; 1465 in in dry-pretreated, low humidity & ABA treated; and 460 in ABA-pretreated, low humidity treated (**Dataset S3**). For each pair of assays, only one member of which caused stomatal closure (non-grey circle in each case), transcripts in each dataset were compared. Environmental conditions and sensitisation status pertaining to each assay are as shown.

**E-F.** Venn comparisons of transcripts associated with stomatal closure, comparing transcripts up-regulated (**E**) and down-regulated (**F**) for each dataset identified in (**A-D**). The sum of each Venn category between (**E**) and (**F**) corresponds to the equivalent category in **Fig. 4*A***.

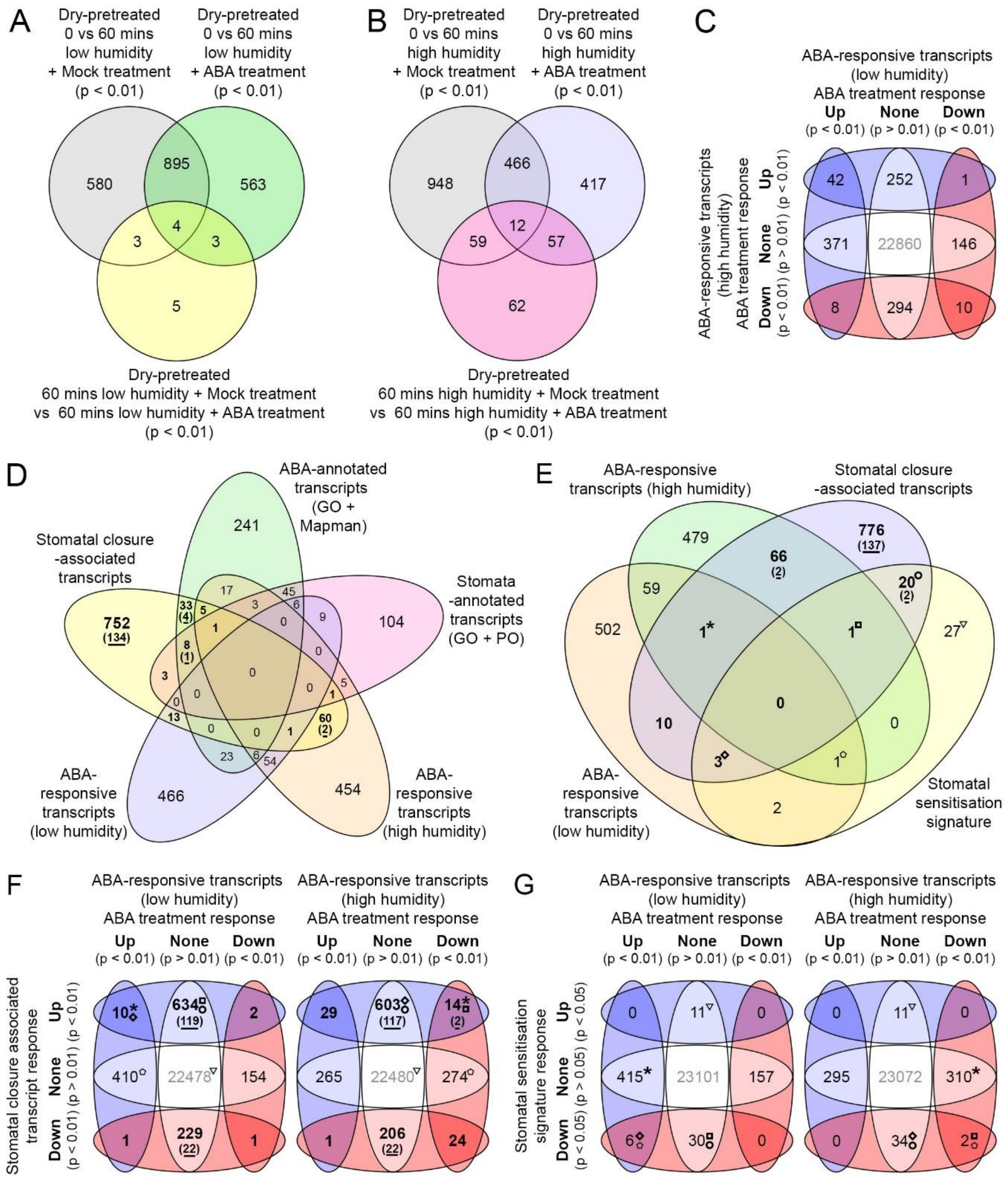


**Fig. S6.** Humidity-dependent ABA regulation of stomatal closure-associated and stomatal sensitisation signature transcripts.

**A,B.** Venn comparison between transcript changes in fronds after 60 minutes mock and ABA treatment under low humidity (**A**) and high humidity (**B**). Transcripts which showed a significant change in abundance ($p < 0.01$) in ABA-treated fronds but not mock-treated fronds, or which showed a significant difference in abundance ($p < 0.01$) between mock and ABA-treated fronds after 60 minutes were classified as ABA-responsive.

**C.** Venn comparison of directional changes in level of ABA-responsive transcripts (**A,B**) after ABA treatment under low and high humidity. The majority of shared transcripts show similar directional responses to ABA treatment under both humidity conditions.

**D.** Venn comparison between stomatal closure-associated transcripts (bold, see **Fig. 4*A***), all transcripts with annotations relating to ABA and stomata functions, and all ABA-responsive transcripts under low or high humidity (**A,B**). Underlined numbers given in brackets denote closure signature transcripts from within the closure-associated transcript dataset.

**E.** Venn comparison between stomatal closure-associated transcripts (bold), stomatal sensitisation signature transcripts (see **Fig. 3*A***) and all ABA-responsive transcripts under low or high humidity (**A,B**). Underlined numbers given in brackets denote closure signature transcripts from within the closure-associated transcript dataset. Superscript symbols denote categories of interest. One closure-associated transcript is ABA-responsive under both low and high humidity (asterisk), as is a single transcript specific to the sensitisation signature (pentagon). Three transcripts common to both the closure-associated and sensitisation datasets are ABA-responsive under low humidity (diamond) while a fourth is ABA-responsive under high humidity (square). The AtPDR12 homolog identified within the sensitisation signature (**Fig. S4**, open circle) is also a closure-associated transcript, but neither this nor the AtMRP4 homolog (**Fig. S4**, open triangle) are ABA-responsive.

**F,G.** Venn comparison between the abundance changes of either stomatal closure-associated transcripts (**F**, bold) or stomatal sensitisation signature transcripts (**G**) during ABA treatment under low and high humidity. Underlined numbers given in brackets (**F**) denote closure signature transcripts within the closure-associated transcript dataset. Superscript symbols within a response category correspond to the categories identified in (**E**). The single closure-associated transcript responsive to ABA under both low and high humidity (asterisk) increases abundance during closure and is similarly regulated by ABA under low humidity. Under high humidity, however, abundance is decreased after ABA treatment (**F**). Conversely, the single sensitisation signature transcript that is responsive to ABA under both humidity conditions (pentagon) shows opposite abundance changes between sensitisation and ABA treatment under low humidity but similar changes between sensitisation and ABA treatment under high humidity (**G**). The abundance of the AtPDR12 homolog (open circle) responds oppositely to closure and sensitisation (**F,G**).

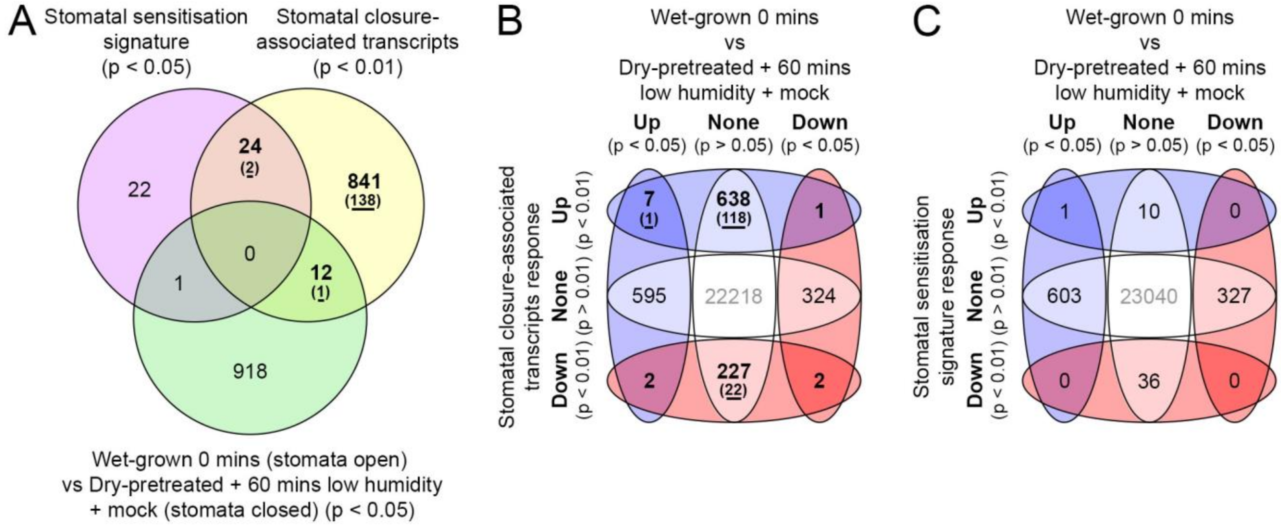


**Fig. S7.** Closure-associated transcripts down-regulated by sensitisation return to their original levels during or after stomatal closure.

**A.** Venn comparison between transcripts where abundance is significantly altered ($p < 0.05$) by sensitisation of wet-grown fronds (see **Fig. 3**), significantly altered ($p < 0.01$) by stomatal closure in sensitised (dry-pretreated) fronds (bold, see **Fig. 4*A***), and significantly different ($p < 0.05$) between wet-grown (open stomata) and dry-pretreated fronds with closing stomata (60 minutes low humidity stimulus). Underlined numbers given in brackets denote closure signature transcripts from within the closure-associated transcript dataset. All 24 transcripts significantly and antagonistically regulated by both sensitisation and closure (see **Fig. 5**) show no significant difference ($p > 0.05$) in abundance prior to sensitisation and after closure. Only a single sensitisation transcript (TRINITY_DN1054_c0_g1) and 12 closure-associated transcripts (including one closure signature transcript) showed a significant difference between levels in sensitised fronds after stomata closure and original levels in wet-grown fronds prior to sensitisation (**Dataset S3**).

**B,C.** Venn comparison between the responses of transcripts with significantly altered abundance between wet-grown fronds (open stomata) and dry-pretreated fronds after stomatal closure, and the responses of closure-associated (**B**) and sensitisation signature (**C**) transcripts during closure. Seven of the 12 closure-associated transcripts identified in (**A**), including the closure signature transcript, showed increased abundance during closure and remained at greater abundance than in wet-grown fronds. The single sensitisation transcript identified in (**A**) increased in abundance in response to sensitisation and remained at greater abundance than in wet-grown fronds.
